# Guided proposals for efficient weighted stochastic simulation

**DOI:** 10.1101/641894

**Authors:** Colin S. Gillespie, Andrew Golightly

## Abstract

Rare event probabilities play an important role in the understanding of the behaviour of biochemical systems. Due to the intractability of the most natural Markov jump process representation of a system of interest, rare event probabilities are typically estimated using importance sampling. While the resulting algorithm is reasonably well developed, the problem of choosing a suitable importance density is far from straightforward. We therefore leverage recent developments on simulation of conditioned jump processes to propose an importance density that is simple to implement and requires no tuning. Our results demonstrate superior performance over some existing approaches.

## 1 Introduction

Simulating discrete biochemical networks via stochastic simulations in recent years has become routine. The default algorithm for many practitioners was developed over four decades ago by Gillespie (1977). This algorithm simulates a single stochastic realisation from the Markov jump process (MJP) representation of a system of interest. Typically, multiple realisations are generated in order to estimate state probabilities. Given the intractability of the MJP, the importance of the stochastic simulation algorithm (SSA) cannot be understated.

Multiple improvements have been proposed to the *default*. For example, Gibson and Bruck (2000) use efficient data structures, whereas other authors McCollum et al. (2006); Slepoy et al. (2008) have leveraged clever sorting strategies. It is also possible to speed up a single stochastic simulation by grouping sets of reactions together and simulating on a multi-core CPU(Gillespie, 2012).

Despite these improvements in computational efficiency, the problem of estimating probabilities of rare events remains challenging. While an ensemble simulation approach can in principle be routinely applied, the number of simulations required to attain a certain level of accuracy can be computationally prohibitive.

The weighted stochastic simulation algorithm (wSSA) was developed by Kuwahara and Mura (2008) to alleviate the computational burden associated with calculating rare event probabilities, that is, estimating *p*(*x*_0_, *x*′ *t*′), the probability of reaching *x*′ before time *t*′, given *x*_0_ at time *t* = 0. The general idea is to carefully *weight* or bias key reactions, which will reduce the total number of simulations required (relative to a vanilla Monte Carlo approach based on the SSA) to give an estimator that attains a given level of accuracy. As an aside, while these simulations can be used to estimate the probability of interest, they can not be regarded as valid realisations from the system of interest. The wSSA can be seen as an importance sampler, and this connection was made explicitly by Gillespie et al. (2009), who further extended the wSSA by proposing to select suitable biasing parameters by *minimising* the variance of the resulting importance sampling estimator. The need for state-dependent biasing parameters was discussed by Roh et al. (2010) who proposed to partition each reaction depending on whether it should be encouraged or not, and specified state-dependent propensity functions for each reaction group. An automated approach to biasing parameter selection was presented by Daigle et al. (2011) with a subsequent extension to state-dependent bias in Roh et al. (2011). In the latter two contributions, a cross entropy method is used to continually refine biasing parameters. Consequently, previous work in this area either requires the specification of additional tuning parameters or additional simulations.

Our novel approach is the specification of an importance density that is simple to implement and requires no additional tuning. We leverage recent developments on simulation of conditioned jump processes to derive a propensity function that takes into account the rare event of interest. We then use this propensity inside the SSA to generate trajectories that are guided towards the rare event *x*′, thereby increasing the likelihood of hitting *x*′ before time *t*′. We find that the proposed approach outperforms competing methods in a variety of settings. Limitations are also discussed.

The remainder of this article is organised as follows. In Section 2, we discuss some recent developments around the wSSA algorithms. Section 3 will consider a simple two-state model, where it is possible to derive analytical expressions for the optimal weights. Following these insights, we will derive our new method in 4 and compare its performance with other methods in Section 5 using three examples of increasing complexity.

## 2 The wSSA Algorithm

Consider a reaction network involving *u* species 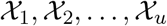 and *v* reaction channels 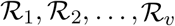 such that

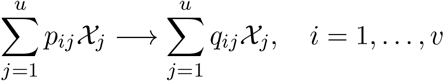

where the *p_ij_* and *q_ij_* are non-negative integers known as stoichiometric coefficients. Let *X_j,t_* denote the (discrete) number of species 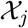 at time *t*, and let *X_t_* be the *u*-vector *X_t_* = (*X*_1, *t*_, *X*_2,*t*_, …, *X_u,t_*)^*T*^. The time evolution of *X_t_* is most naturally described by a Markov jump process (MJP), so that for an infinitesimal time increment *dt* and a propensity function (or instantaneous hazard) *h_i_*(*x_t_*), the probability of a type *i* reaction occurring in the time interval (*t*, *t* + *dt*] is *h_i_*(*x_t_*)*dt*. Under the standard assumption of mass action kinetics, *h_i_* is proportional to a product of binomial coefficients. Specifically

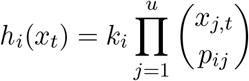

and we suppress explicit dependence of *h_i_* on the rate constant *k_i_*. Values for *k* = (*k*_1_, *k*_2_, …, *k_v_*)^*T*^, the initial system state *X*_0_ = *x*_0_ and the *u* × *v* stoichiometry matrix *S* whose (*i*, *j*)th element is given by *q_ji_* – *p_ji_*, complete specification of the Markov process. Despite the intractability of the probability mass function governing the state of the system at any time *t* (as satisfies the chemical master equation), generating exact realisations of the MJP is straightforward via a technique known in this context as the *stochastic simulation algorithm* (SSA) (Gillespie, 1977). In brief, if the current time and state of the system are *t* and *X_t_* respectively, then the time to the next event will be exponential with rate parameter

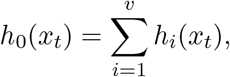

and the event will be a reaction of type 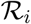 with probability *h_i_*(*x_t_*)/*h*_0_(*x_t_*) independently of the inter-event time.

We suppose that interest lies in *p*(*x*_0_, *x*′; *t*′), that is, the probability that the system starting at the initial value *x*_0_ at time 0 will first reach the state *x*′ at some time *t* in the interval (0, *t*′]. This probability can be written as the expectation

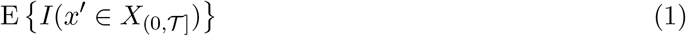

where the indicator function 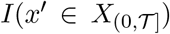 takes the value 1 if 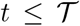 and 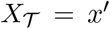 at some stopping time 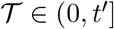. It takes the value 0 otherwise. Note that the expectation is with respect to the probability law of the MJP over the interval (0, *t*′], denoted *p*(*x*_(0,*t*′]_|*x*_0_). Then, a simple Monte Carlo procedure for (unbiasedly) estimating the expectation in (1) is to generate (say) *N* independent draws *x*^(1)^, …, *x*^(*N*)^ of 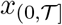 via the SSA and set

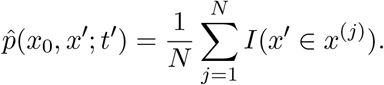

However, it is well known that when dealing with rare events, the corresponding estimator will have high variance (for a given *N*), since relatively few draws of the MJP will attain the condition for setting the indicator equal to 1 (see Kuwahara and Mura, 2008).

The weighted SSA (wSSA) constructs an *importance sampling* estimate of *p*(*x*_0_, *x*′; *t*′). A key ingredient of this approach is a suitable *importance density q*(*x*_(0, *t*′]_|*x*_0_). Given *N* independent draws *x*^(1)^, …, *x*^(*N*)^ of 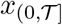 from 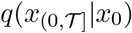, the importance sampling estimate is

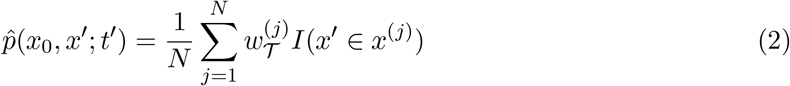

where the *weight* function is

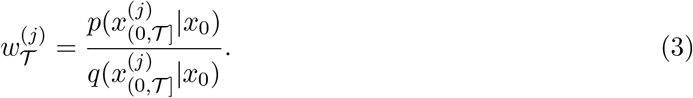

The importance density should be chosen so that samples drawn from it are “more likely” to meet the attainment condition. Hence, the choice of importance density is crucial for devising an estimator with low variance. In what follows, we take 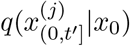 to be the probability law associated with an MJP whose propensity function is 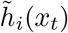 for reaction *i* = 1, …, *v*. In this case, the weight function can be explicitly calculated as follows. We let *r_j_* denote the number of reaction events of type 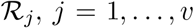, and define 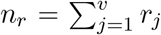 as the total number of reaction events over the interval 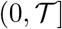. Reaction times (assumed to be in increasing order) and types are denoted by (*t_i_*, *ν_i_*), *i* = 1, …, *n_r_*, *ν_i_* ∈ {1, …, *v*} and we take *t*_0_ = 0 and 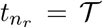. The *complete-data likelihood* over 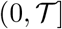 is

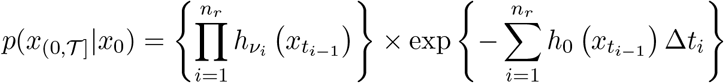

where Δ*t_i_* = *t_i_* – *t*_*i*–1_ is the dwell time preceding the *i*th reaction firing (see Wilkinson, 2012). An expression for 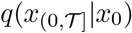 is obtained similarly. Hence the weight function is

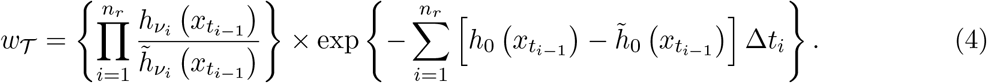

Computation of an estimate 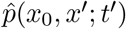 is then straightforward by performing *N* independent runs of the SSA with *h_i_*(*x_t_*) replaced by 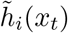. The number *M_N_* of runs that reach state *x*′ prior to time *t*′ is obtained, and each such run is assigned a weight using (4). The weights are then used as in (2) to estimate *p*(*x*_0_, *x*′; *t*′).

It remains that we specify a suitable form of 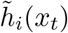 to be used in the importance sampler. Previous approaches (Kuwahara and Mura, 2008; Gillespie et al., 2009) have used *h*_0_(*x_t_*) to update the inter-event times and 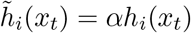 to update the reaction events, for some tuning parameter *α*. Roh et al. (2011) use *αh_i_*(*x_t_*) to update both the inter-event times and reaction events. In the following section we motivate the need for the scaling parameter *α* to be time-dependent before presenting our novel approach in Section 4.

## 3 Discrete two state model

Consider a simple, *discrete-time*, two state model. Suppose *X* can only take that the values 0 or 1. The probability of remaining in state 0 is *p*_0_. The probability of moving from state 0 to 1 is *p*_1_ = 1 – *p*_0_. The state *X* = 1, is an absorbing state.

The system commences with *X* = 0, at time *t* = 0. Hence, the probability that the system will first reach the state 1 at some time *t* in the interval (0, *t*′] is

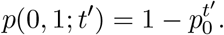

Suppose we wish to encourage absorption, i.e. reaching state 1 by time *t*′. An importance sampling strategy is to weight the probability of a transition from 0 to 1, by *α*.

Hence at time *t*, assuming we have not reached state 1, we draw *X_t_* according to the probability mass function with

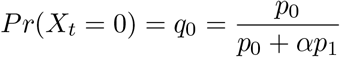

and

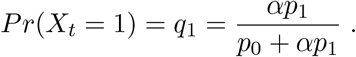

Then, the associated weight should be multiplied by one of

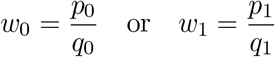

depending on the transition type. The expected weight at time *t*′ is given by

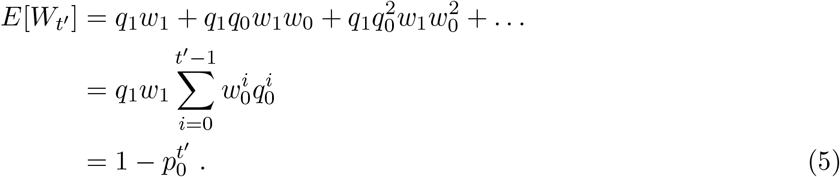

Hence, the importance weights can be used to unbiasedly estimate *p*(0, 1; *t*′) for any *α* > 0. However, as Gillespie et al. (2009) showed, the efficiency of the method can vary dramatically with the choice of *α*. To optimise the choice *α*, we minimise the variance of the estimate. For this model, we can derive an analytical expression for the second moment, namely

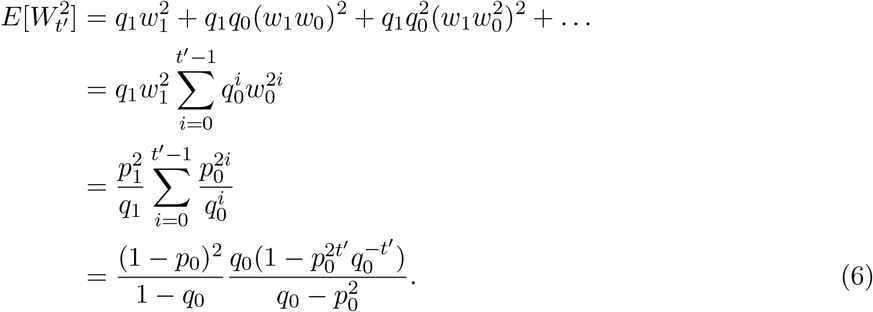

To obtain an optimal value of *α*, we need to minimise

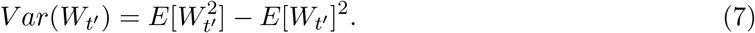

Since *E*[*W_t′_*] does not depend on *α*, minimising *V ar*(*W*_*t*′_) is equivalent to minimising 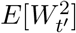. The crucial insight from equation 6, is that 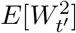 explicitly depends on *t*′. This implies that the optimal value of *α* varies over time. This highlights that the methods proposed by Kuwahara and Mura (2008); Gillespie et al. (2009); Roh et al. (2010) are not optimal, since they assume a fixed value of *α* for all times *t* ∈ [0, *t*′).

For the case when *t*′ → ∞, the expectation simplifies to

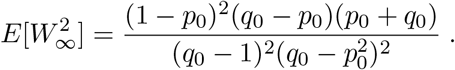

This is minimised by *q*_0_ = *p*_0_, i.e. setting *α* = 1. This makes intuitive sense. Essentially as *t*′ increases, the probability of hitting the target approaches 1. Hence, every simulation will eventually hit the target and so adjusting the probabilities is not needed.

For the general case, we can minimise 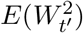 by solving

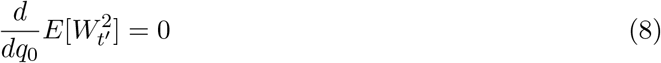

for *q*_0_. After some differentiation we have that

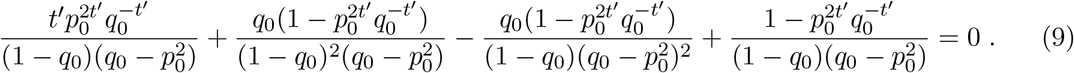

Although equation (9) cannot be solved analytically, a standard numerical solver can be used to obtain *q*_0_ for a given value of *t*′. Suppose that *p*_0_ = 0.9 and interest lies in hitting *X* = 1 by time *t*′ = 5. The optimal value of *α* (and therefore *q*_0_) can be calculated for *t* = 0, 1 …, 4, under the assumption that for each *t*, we have not yet reached the absorbing state. For example, if at *t* = 4 we have not yet reached state 1, the optimal *α* is the value that minimises 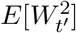 with *t*′ = 5 – *t* = 1. Figure 1 shows that as *t* increases, and approaches *t*′ = 5, the optimal value of *α* increases accordingly. When *t* = 0, *α* = 2.2. This corresponds to *q*_1_ = 0.20 (*p*_1_ = 0.1). However, when *t* = 4, i.e. the last possible time at which *X_t_* can be pushed towards state 1, *α* = 13.51 and *q*_1_ = 0.60.

**Figure 1:**
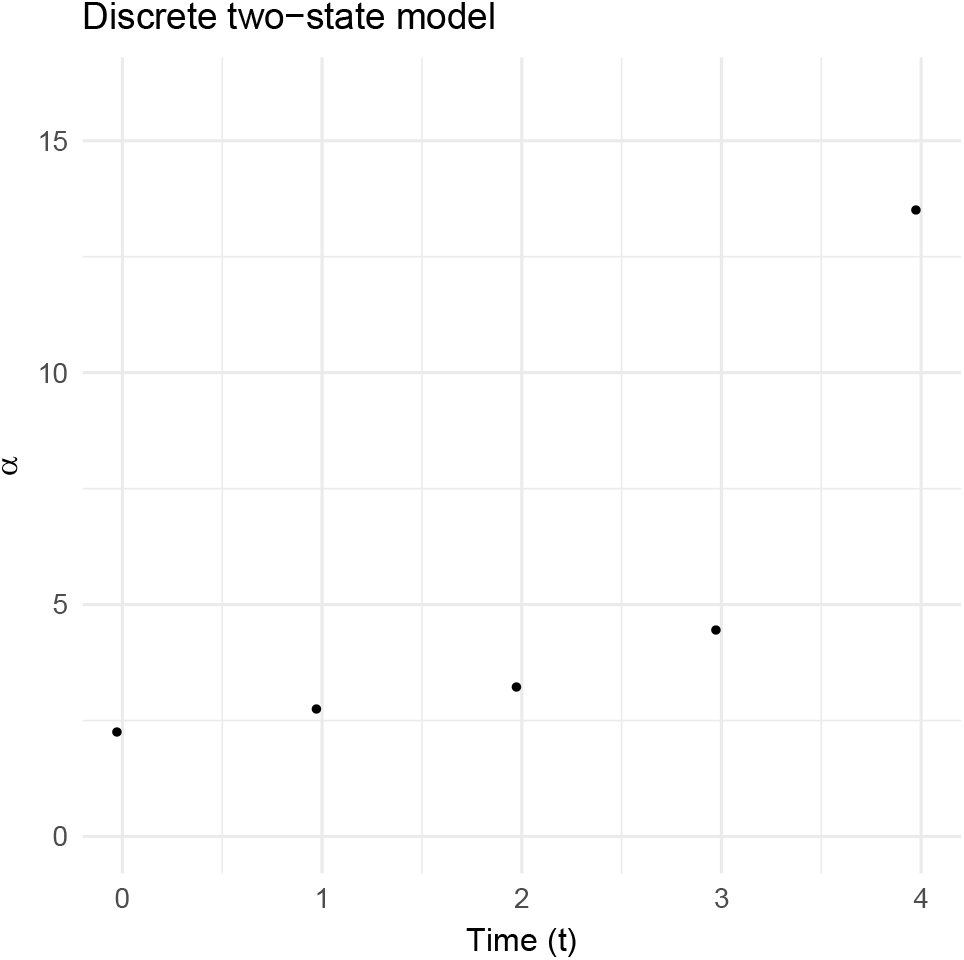
Optimal values of *α* against *t* for the simple two-state model, with *p*_0_ = 0.9 and *t*′ = 1.

## 4 Guided wSSA

To obtain an importance density 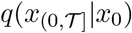, a suitable form of 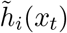 (for use in the SSA) is required. We appeal to the recent literature on the sampling of jump processes that are conditioned to match some value at a given future time (Golightly and Wilkinson, 2015). Essentially, we seek a tractable approximation of the expected number of reaction events in an interval of the form (*t*, *t*′]. The resulting expression is used as the propensity function in the SSA, giving draws from the corresponding 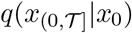.

In what follows we assume that

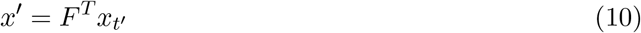

for some constant matrix *F* of dimension *u* × *u_o_*, with *u_o_* denoting the number of components of *x*′. Hence, we allow for the rare event of interest to depend on a linear combination of the components of the MJP. For simplicity, consider the generation of a single draw from the importance density 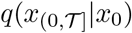. Suppose that we have simulated as far as time 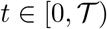. Let Δ*R_t_* denote the number of reaction events over the time *t*′ – *t* = Δ*t*. We approximate Δ*R_t_* by assuming a constant reaction hazard over Δ*t*. A Gaussian approximation to the corresponding Poisson distribution then gives

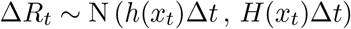

where *H*(*x_t_*) = diag{*h*(*x_t_*)}. By linearity of (10) and using *X_t_* = *S*Δ*R_t_*, we have that

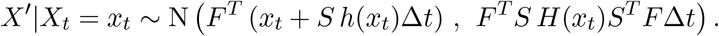

Hence, the joint distribution of Δ*R_t_* and *X*′ (conditional on *x_t_*) can be obtained approximately as

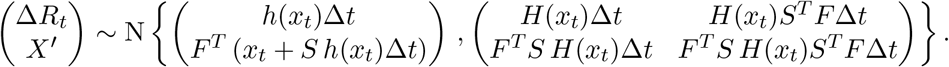

Taking the expectation of Δ*R_t_*|*X*′ = *x*′ and dividing the resulting expression by Δ*t* gives an approximate conditioned hazard as

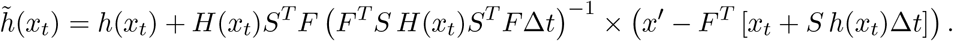

### Algorithm 1 Guided wSSA

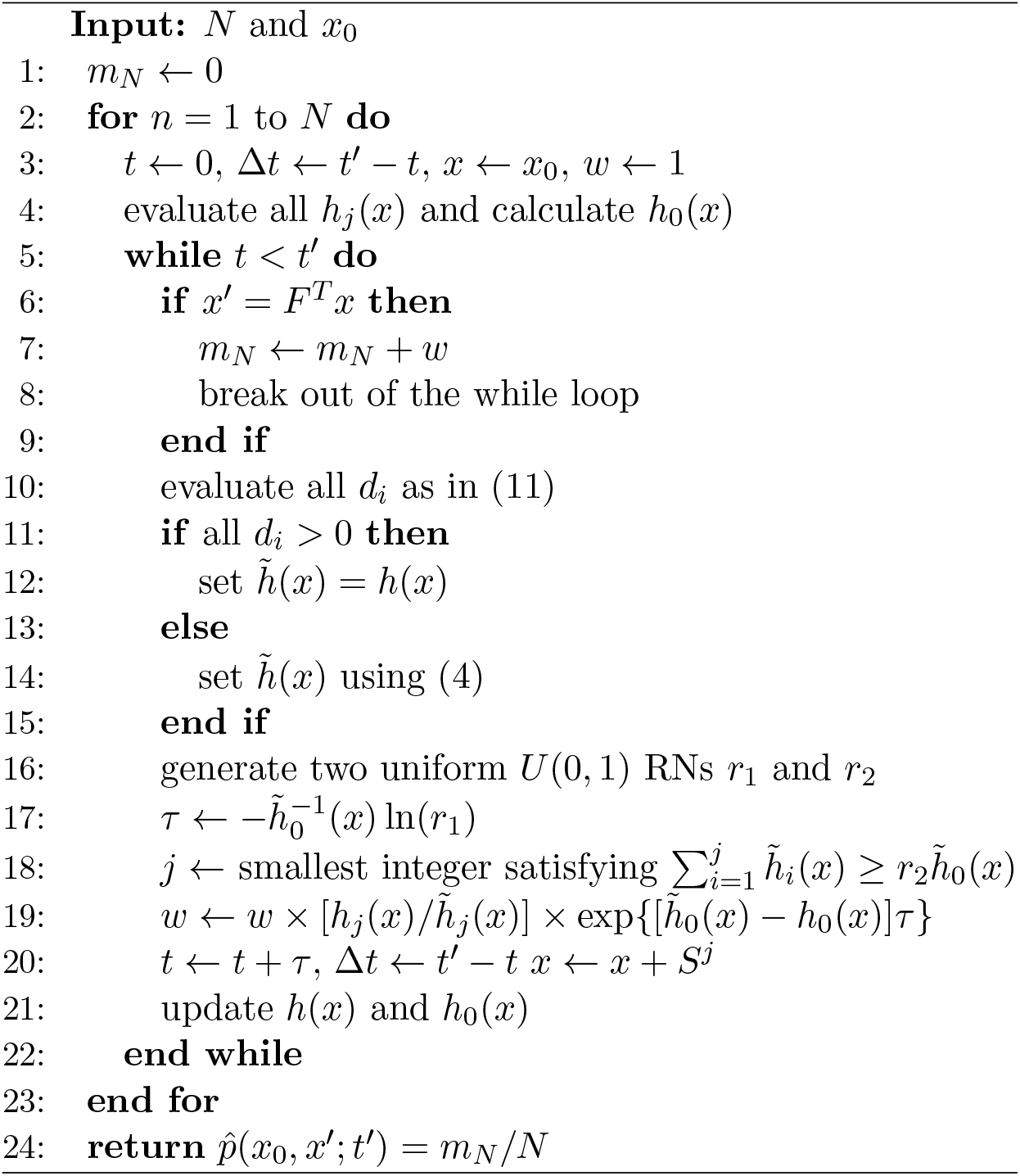

Using 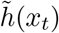 inside the SSA requires calculation of the conditioned hazard function in (4) after every reaction event. The cost of this calculation will be dictated by the dimension, *u_o_*, of *x*′, given that a *u_o_* × *u_o_* matrix must be inverted. We anticipate that it will often be the case that *u_o_* ≪ *u*, for example when one gene is of primary interest in a larger system. Nevertheless, to avoid unnecessary computational cost, we suggest that at time *t*, a *pre-simulation check* is performed. We have approximately that

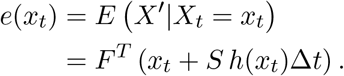

Hence, we compare *E* (*X*′|*X_t_* = *x_t_*) to *x*′ by computing

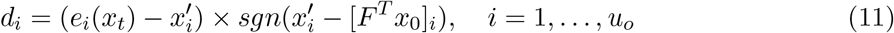

where, for example, [*F^T^ x*_0_]_*i*_ denotes the *i*th component of *F^T^ x*_0_. Hence if all values *d_i_* are positive, no additional push towards *x*′ is necessary and we take 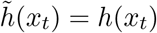, thus avoiding calculation of the inverse of a *u_o_* × *u_o_* matrix. We note that 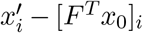 need only be computed once, and that the calculation of *e*(*x_t_*) should be inexpensive, since it solves simple operations of the pre-specified quantities *S* and *F*, and the current hazard *h*(*x_t_*). Algorithm 1 gives the full wSSA (denoted “guided wSSA”) scheme based on the conditioned hazard function described above. Lines 10–15 describe the pre-simulation check.

The construction of the conditioned hazard is based on an assumption that the hazard function is constant over diminishing time intervals (*t*, *t*′] and that the number of reactions over this interval is approximately Gaussian. The performance of the construct is therefore likely to be reduced if applied over time horizons during which the reaction hazards vary substantially. Additionally, the Guassian assumption is likely to be unreasonable in low count scenarios, although we note that in such scenarios, simply using the SSA is likely to computationally feasible. In what follows we investigate the performance of the proposed approach in three examples.

## 5 Examples

We consider three applications of increasing complexity. We compare and contrast the guided wSSA with vanilla use of the SSA, the wSSA of Kuwahara and Mura (2008) and the state-dependent wSSA of Roh et al. (2010). Following Gillespie et al. (2009), we compare the accuracy of each method by computing the sample variance of the unnormalised importance weights. That is the sample variance of

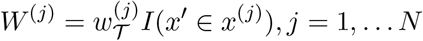

where 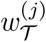 is given by (3). We denote the resulting quantity by *σ*^2^. All methods were coded in R and run on were run on a desktop computer with an Intel Core i7-4770 processor at 3.40GHz

### 5.1 Single species production and degradation

Our first example is taken from Kuwahara and Mura (2008) (and see also Gillespie et al. (2009) and Roh et al. (2010)). The system contains two reactions

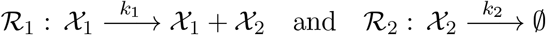

where *k*_1_ = 1 and *k*_2_ = 0.025. The initial state is *x*_0_ = (1,40)^*T*^ we wish to estimate *p*(*x*_0_, *F*^*T*^ *x*_*t*′_; *t*′) with *t*′ = 100, *F^T^* = (0,1) and *x*′ = *F*^*T*^ *x*_*t*′_ = 80. That is, the probability that *X*_2,*t*_ reaches *x*′ = 80 before time *t*′ = 100 given an initial state of *x*_0_ = (1,40)^*T*^ at *t* = 0. This is a well known model and the steady state population of *X*_2,*t*_ follows a Poisson distribution with parameter λ = *k*_1_*x*_1,*t*_/*k*_2_ where *x*_1,*t*_ = 1 for all times *t*. Since the initial state *X*_2,0_ is set at the equilibrium value, *X*_2,*t*_ fluctuates around 40 with a standard deviation of 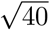. Hence, *x*′ = 80 is the upper 10^−6^% quantile of the equilibrium distribution.

Figure 2 shows 3 trajectories that meet the attainment condition, generated using the guided wSSA (Algorithm 1). For this example, the pre-simulation check is performed at time *t* by computing *e*(*x_t_*) – *x*′ where

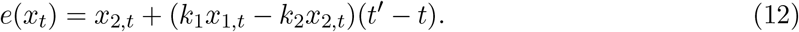

**Figure 2:**
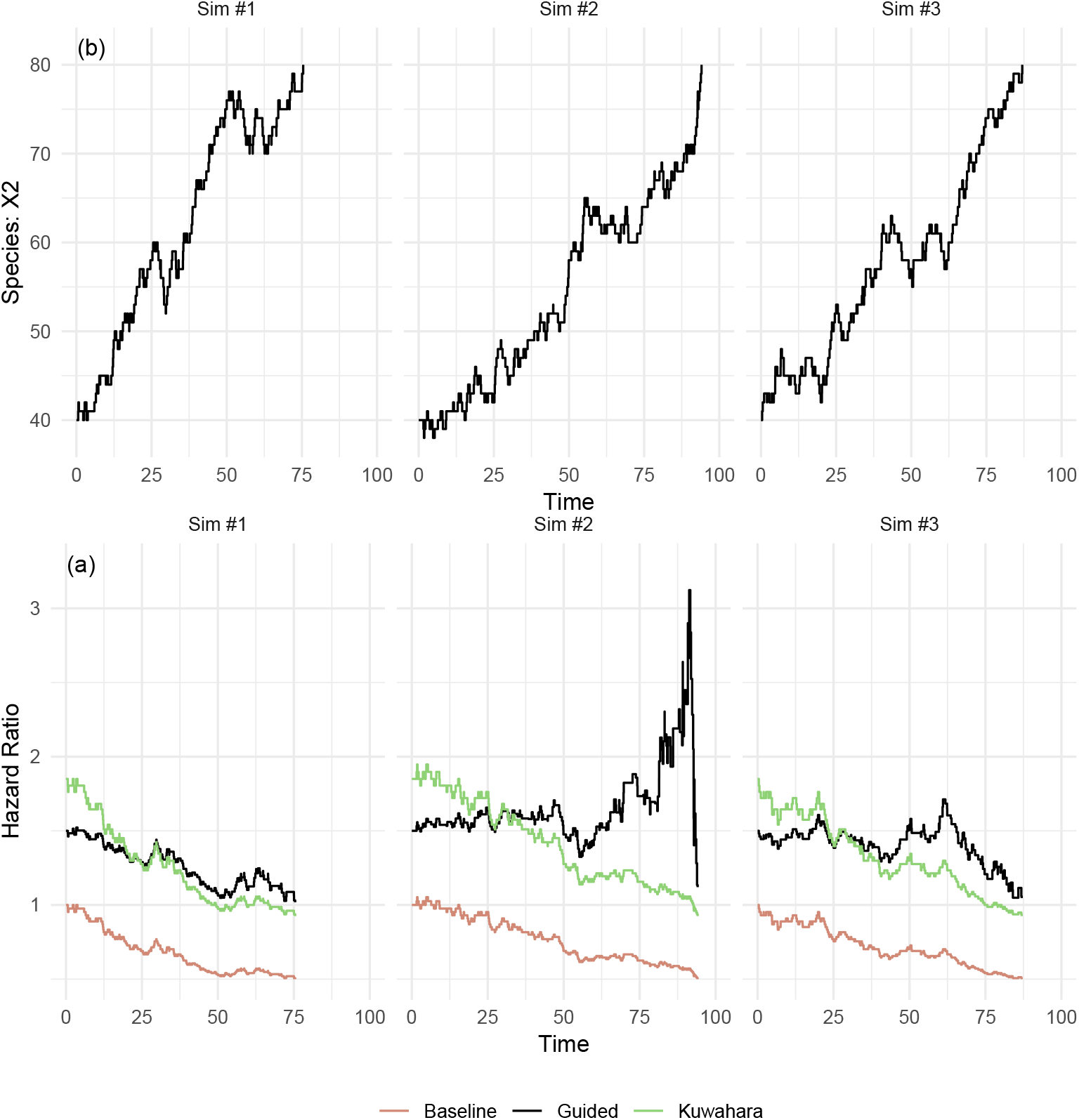
Top panel (a): 3 trajectories using the guided SSA that attain *x*′ = 80 by *t*′ = 100. Bottom panel (b): ratios 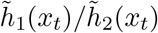 against time *t* (where *x_t_* is given in the top panel) under the guided wSSA (black) and Kuwahara/Mura approach (green). The ratio of unaltered hazards *h*_1_(*x_t_*)/*h*_2_(*x_t_*) is also shown (red line).

If *x*′ – *e*(*x_t_*) < 0, we take 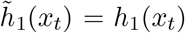 and 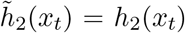. If *x*′ – *e*(*x_t_*) > 0, we use (4) to obtain

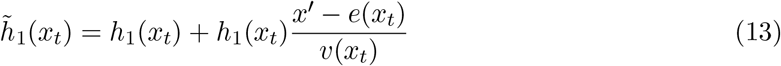

and

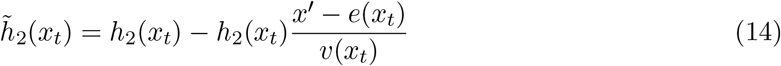

where *v*(*x_t_*) = (*k*_1_ *x*_1,*t*_ + *k*_2_*x*_2,*t*_)(*t*′ – *t*). In this case, the propensity of reaction 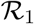 is increased and the propensity of reaction 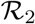 is decreased.

Based on the trajectories in Figure 2, we computed the ratio 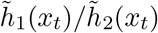. We additionally computed the same quantity under the wSSA method of Kuwahara and Mura (2008), given by

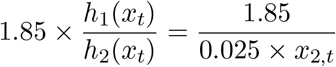

corresponding to 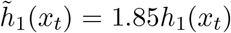 and 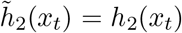. Hence the Kuwahara/Mura approach is to bias reaction 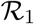 by increasing its propensity. We also display the ratio of unaltered hazard functions

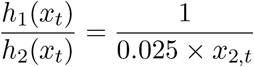

which we refer to as the baseline. We see that the propensity ratios under the baseline and Kuwahara/Mura approach exhibit a decreasing trend, and although the latter is able to increase the propensity of reaction 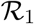, neither method is able to adapt to changes in *x_t_*. The guided wSSA, on the other hand, is able to adjust the relative propensity depending on the behaviour of *x_t_*. For example, in simulation 2 (middle column of Figure 2), the trajectory appears to be fairly constant between the values 60–65 between times 55–75. The guided wSSA therefore gives a sharp increase in relative propensity in order to meet the attainment condition.

We ran guided wSSA and the wSSA of Kuwahara/Mura for *N* = 10^5^ trajectories and computed the sample variance *σ*^2^ of the importance weights. For comparison, we also ran the vanilla SSA approach (using the unaltered relative propensity function given above), which required *N* = 10^8^ to give a reasonable number of trajectories that meet the attainment condition. This gave variances of 1.1 × 10^−12^ for guided wSSA, 1.5 × 10^−10^ for the Kuwahara and Mura approach and 2.7 × 10^−7^ for the vanilla SSA approach. Hence, guided wSSA outperforms wSSA by two orders of magnitude and SSA by five orders of magnitude.

### 5.2 Two-state conformational transition

The following simple reaction network, initially described by Kuwahara and Mura (2008), models two conformational isomers, that is, isomers that can be inter-converted by rotation about single bonds. This model has two reactions

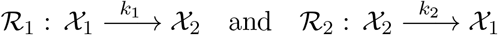

where *k*_1_ = 0.12 and *k*_2_ = 1. The initial state is set to *x*_0_ = (100, 0)^*T*^. For this particular system we wish to estimate *p*(*x*_0_, *F*^*T*^*x_t′_*; *t*′) with *t*′ = 10, *F^T^* = (0,1) and *x*′ = *F*^*T*^*x_t′_* = 30. That is, the probability that given *x*_0_ = (100, 0)^*T*^, the value of *x*_2,*t*_ reaches *x*′ = 30 before time *t*′ = 10. The steady state population of *X*_2,*t*_ is approximately 11, so the rare event of *x*′ = 30 by time *t*′ = 10, is around three times larger than it’s equilibrium value. Since this system is closed, we can numerically calculate this probability to obtain

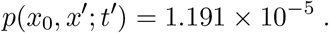

Since the steady state population of *X*_2,*t*_ is less than the target value of 30, we can bias the system by either increasing the rate of reaction *R*_1_ or decreasing the rate of reaction *R*_2_.

The implementation of the guided wSSA follows closely the approach described in Section 5.1. The pre-simulation check calculates *e*(*x_t_*) – *x*′ where *e*(*x_t_*) is given by (12). For the non-trivial case of *x*′ – *e*(*x_t_*) > 0, 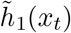 and 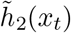 are given by (13) and (14) respectively. We compare the output of the guided wSSA against that of the wSSA described by Kuwahara and Mura (2008) and the state-dependent approach of Roh et al. (2010)). For the former, the first reaction *R*_1_, was encouraged via the biasing parameter *γ*. This corresponds to

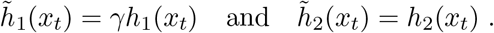

The optimal biasing parameter was estimated Roh et al. (2010) by selecting the value of *γ* that minimised the variance of the estimator of *p*(*x*_0_, *x*′; *t*′). This resulted in *γ* = 1.4. The Roh et al. method encourages reaction 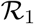 with the state-dependent biasing parameter

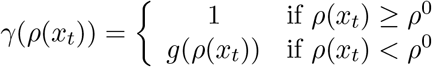

where

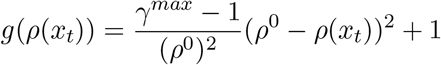

and *ρ*(*x_t_*) = *h*_1_(*x_t_*)/*h*_0_(*x_t_*). As discussed in Roh et al. (2010), *g*(*ρ*(*x_t_*)) has the desirable properties that *g*(0) = *γ^max^*, the maximum biasing value allowed, and *g*(*ρ*^0^) = 1, to avoid over-perturbing the system beyond a relative propensity of *ρ*^0^. For this example, the authors suggest *ρ*^0^ = 0.5 and *γ^max^* = 20.

Figure 3 shows the sample variance *σ*^2^ of the importance weights obtained from four runs of each method (and additionlly, the vanilla SSA approach) using *N* = 10^5^. The guided wSSA and Roh et al. approaches give comparable variances whereas the wSSA of Kuwahara/Mura gives variances that are typically at least two orders of magnitude greater than those exhibited by the former. Simply using the SSA gives importance weights whose variance is around four orders of magnitude greater than the guided approach. We stress the automatic nature of the guided wSSA approach - no tuning parameters are required. For further comparison, we also display results for *t*′ = 30.

**Figure 3:**
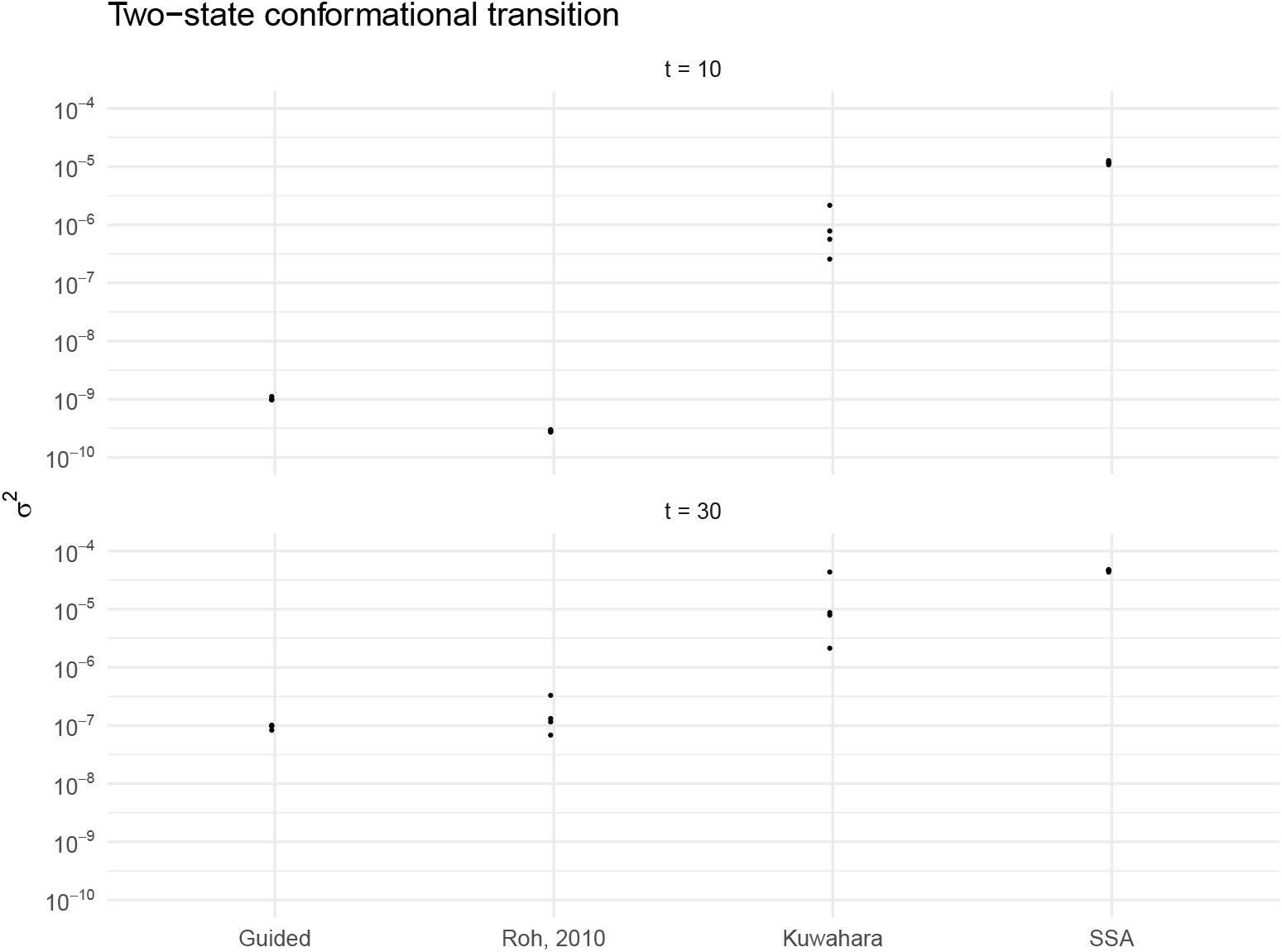
A plot of *σ*^2^ for the two-state conformational model. Each of the four points is calculated using *N* = 10^5^ for the guided wSSA and two competing methods. Upper panel (*t* = 10): The variance associated with estimating *p*((100,0)^*T*^, 30; *t*′ = 10) and lower panel (*t* = 30): *p*((100, 0)^*T*^, 30; *t*′ = 30).

### 5.3 Motility Regulation

In this section we consider a simplified model of a key cellular decision made by the gram-positive bacterium *Bacillus subtilis* (Sonenshein et al., 2002). This decision is whether or not to grow flagella and become motile (Kearns and Losick, 2005). The *B. subtilis* sigma factor *sigD* is key for the regulation of motility. Many of the genes and operons encoding motility-related proteins are governed by *sigD*, and so understanding its regulation is key to understanding the motility decision. The gene for *sigD* is embedded in a large operon containing several other motility-related genes, known as the *fla/che* operon. Transcription of the operon is strongly repressed by the protein *CodY*, which is encoded upstream of *fla/che. CodY* inhibits transcription by binding to the *fla/che* promoter. Since *CodY* is upregulated in good nutrient conditions, this is thought to be a key mechanism for motility regulation. For simplicity we focus here on one gene under the control of *sigD*, that is *hag*, which encodes the protein *flagellin* (or *Hag*), the key building block of the flagella. The reaction network has *u* = 9 species and *v* = 12 reactions and is encoded as follows:

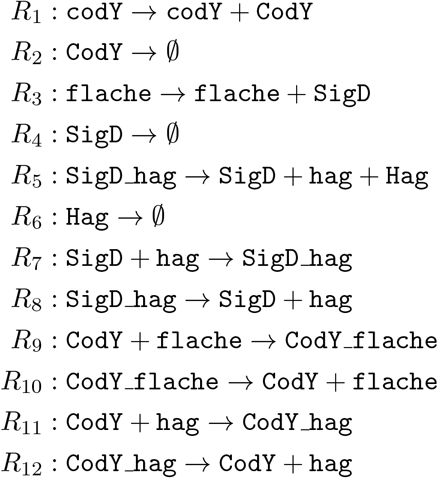

Values of the rate constants are taken to be

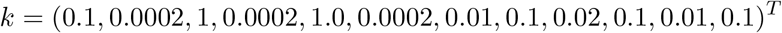

and the initial values of

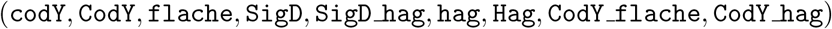

are

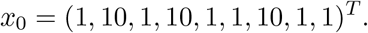

Figure 4 gives ten realisations of the system with these settings. Note that trajectories for each species are inherently discrete. Since the guided wSSA is based on a normal approximation of the MJP, we anticipate that these parameter settings provide a challenging scenario for the proposed methodology.

**Figure 4:**
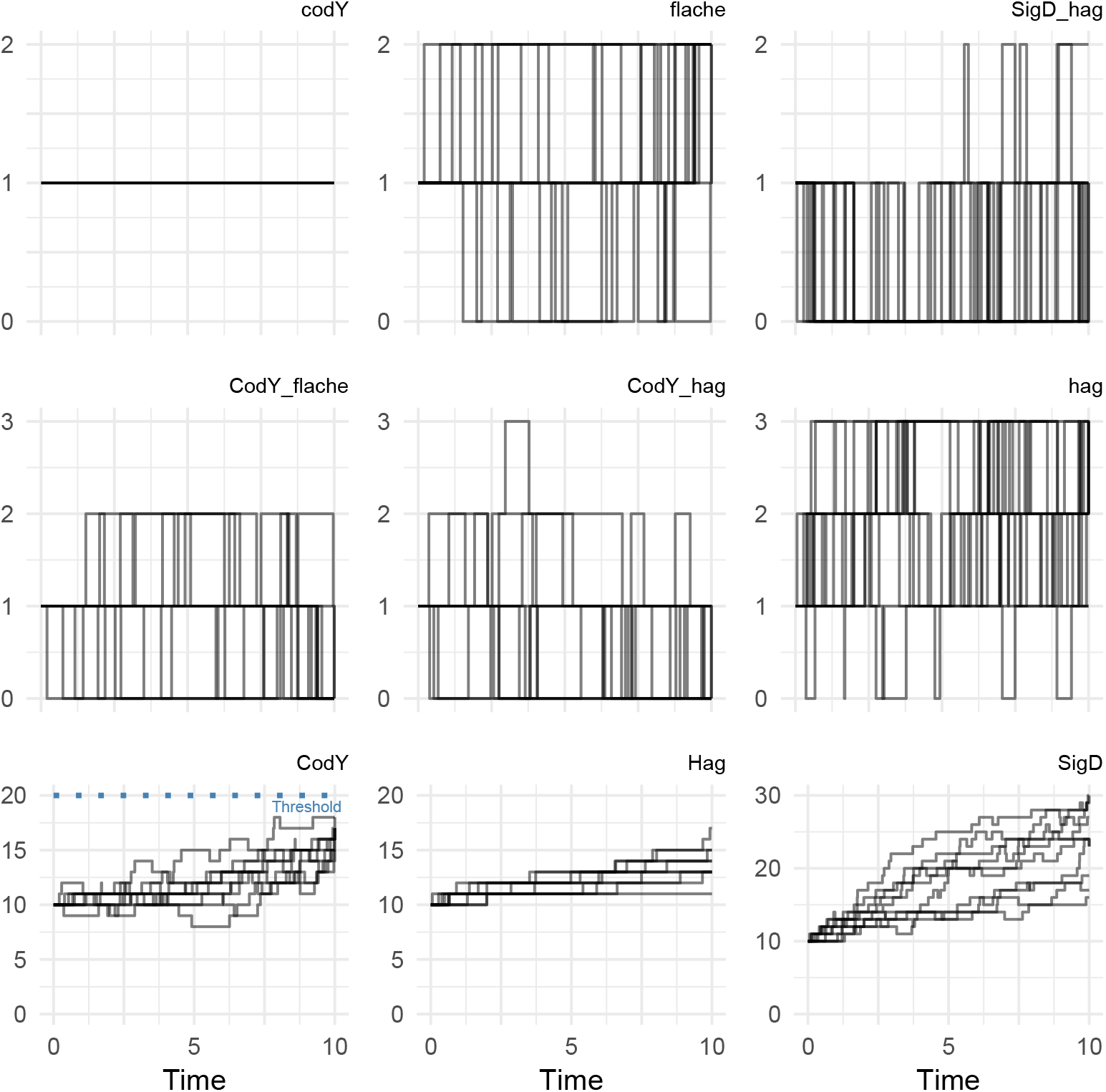
Ten stochastic realisations of the motility regulation model. The rare event of interest is indicated (see bottom left).

Given the importance of *CodY* in motility regulation, we assume that interest lies in the estimation of *p*(*x*_0_, *F^T^x*_*t*′_;*t*′) with *t*′ = 10, *F^T^* = (0,1,0, …, 0) and *x*′ = *F^T^x*_*t*′_ = 20. That is, the probability that given *x*_0_ as above, the value of `CodY` reaches *x*′ = 20 before time *t*′ = 10. Given the size of the reaction network, tuning a weighted SSA approach is likely to be difficult. We therefore compare guided wSSA with a vanilla SSA approach, since neither method requires tuning. For guided wSSA, we used *N* = 10^6^ and for SSA we used *N* = 10^8^, with the latter required to produce a reasonable number of trajectories that met the attainment condition. Our results are reported in Figure 5. The computational cost of guided wSSA versus SSA scales roughly as 2 : 1 for this example. We note however that this is computational effort that is well worth spending, since the guided wSSA gives values of *σ*^2^ (the variance of the importance weights) that are typically three orders of magnitude smaller than when using the SSA.

**Figure 5:**
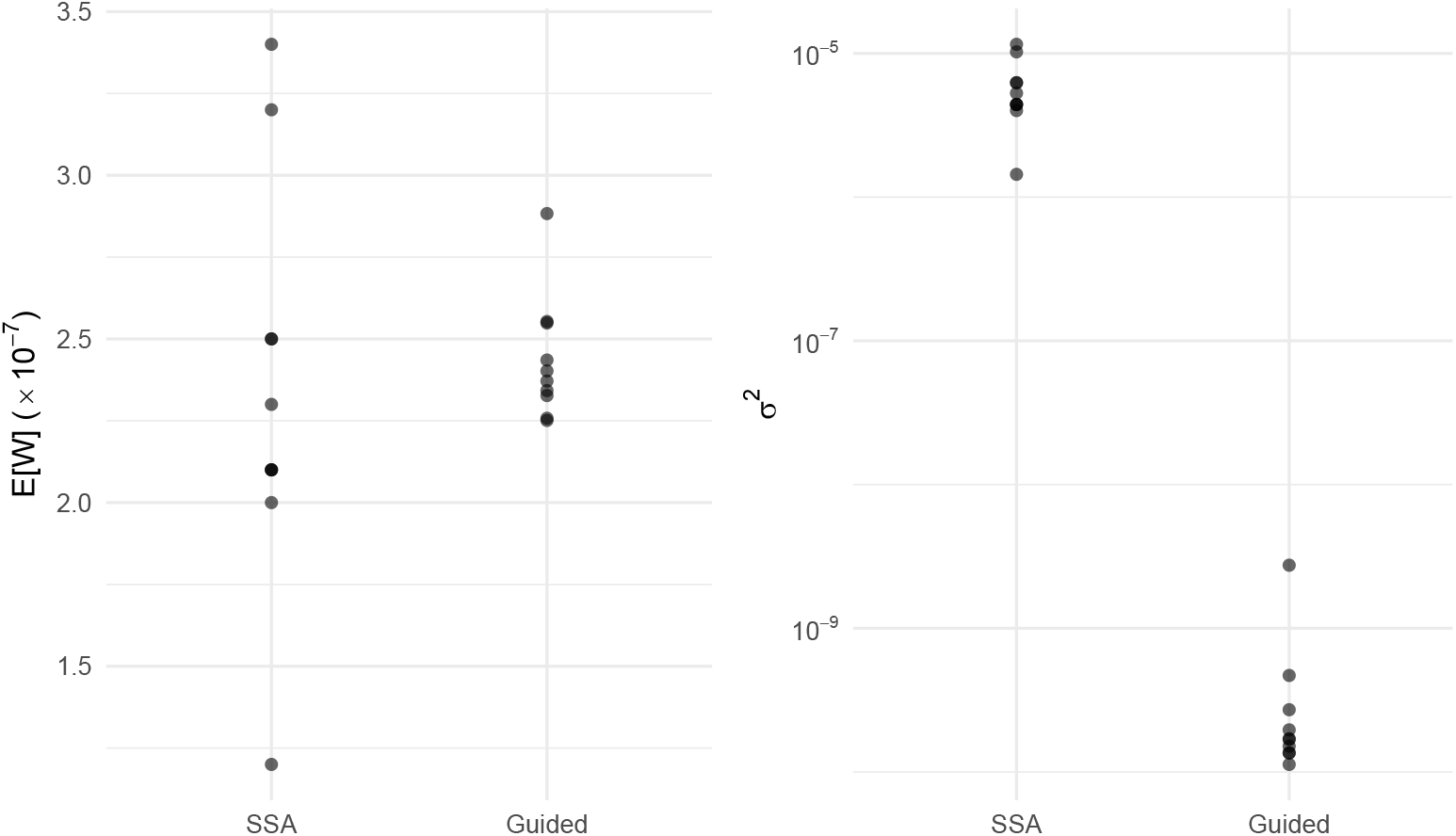
Motility regulation model: expectation (left panel) and variance (right panel) of the importance weights. Each of the eight points is calculated using *N* = 10^6^ for guided wSSA *N* = 10^8^ for SSA.

## 6 Conclusions

Simulation-based approaches to rare event probability estimation can be computationally prohibitive. The weighted stochastic simulation algorithm (wSSA) aims to avoid wasteful simulation by using an importance sampler to push trajectories towards the rare event of interest. The resulting biased trajectories are appropriately weighted, and the weights are averaged to give an unbiased estimator of the rare event probability. Constructing the importance sampler usually requires insight into the underlying reaction system, careful tuning or additional simulations.

In this paper we have introduced an importance sampler that requires no tuning. We have leveraged the tractability of a Gaussian approximation to the Markov jump process to construct a propensity function that is conditioned on the rare event event occurring at a future time. Use of this propensity inside the SSA gives the guided wSSA. Our experiments suggest that the guided wSSA can provide a practical approach to rare event probability estimation, by comprehensively outperforming the unweighted SSA method, and giving comparable results to other wSSA approaches, without the need for laborious determination of suitable tuning parameters. While the Gaussian approximation (on which the conditioned propensity is based) is likely to be unsatisfactory when species exhibit inherently discrete behaviour, our findings suggest that the guided wSSA is relatively robust in such low-count number scenarios. Nevertheless, improving the efficiency of the guided wSSA (e.g. by leveraging more accurate moment-closure based approximations to the MJP) remain the topic of future work.

